# Disentangling Hierarchical and Sequential Computations during Sentence Processing

**DOI:** 10.1101/2022.07.08.499161

**Authors:** Christos-Nikolaos Zacharopoulos, Stanislas Dehaene, Yair Lakretz

## Abstract

Sentences in natural language have a hierarchical structure, that can be described in terms of nested trees. To compose sentence meaning, the human brain needs to link successive words into complex syntactic structures. However, such hierarchical-structure processing could co-exist with a simpler, shallower, and perhaps evolutionarily older mechanism for local, word-by-word sequential processing. Indeed, classic work from psycholinguistics suggests the existence of such non-hierarchical processing, which can interfere with hierarchical processing and lead to sentence-processing errors in humans. However, such interference can arise from two, non mutually exclusive, reasons: interference between words in working memory, or interference between local versus long-distance word-prediction signals. Teasing apart these two possibilities is difficult based on behavioral data alone. Here, we conducted a magnetoencephalography experiment to study hierarchical vs. sequential computations during sentence processing in the human brain. We studied whether the two processes have distinct neural signatures and whether sequential interference observed behaviorally is due to memory-based interference or to competing word-prediction signals. Our results show (1) a large dominance of hierarchical processing in the human brain compared to sequential processing, and (2) neural evidence for interference between words in memory, but no evidence for competing prediction signals. Our study shows that once words enter the language system, computations are dominated by structure-based processing and largely robust to sequential effects; and that even when behavioral interference occurs, it need not indicate the existence of a shallow, local language prediction system.

## Introduction

A central question in linguistics pertains to the nature of linguistic structures and their processing. According to generative linguistics, language processing is enabled via a human-specific ability to mentally transform and represent sequentially incoming words into a hierarchical tree structure (1–4). Others, however, claim that sequential structure has considerable explanatory power and that hierarchical processing is often not involved in language use (5). According to this view, language ability does not rely on an innate competence to process hierarchical structures, and it is rather acquired through a process in which children build up a repertoire of increasingly complex constructions by gradually abstracting from sequential transition probabilities (6, 7).

The ubiquity of hierarchical structures in language has been shown, however, in numerous analyses of linguistic phenomena such as binding (8), question construction (9) language acquisition (10, 11), prosody (12) and phonology (13, 14). Neuroimaging studies have provided further support for the hierarchical view by showing that neural activity in specific regions of the human brain is sensitive to the hierarchical structure of sentences (e.g., 15–22). Together, these studies provide largely incontrovertible evidence for nested tree-like representations in humans, making the hierarchical view currently dominant in the field.

However, hierarchical structure processing could co-exist with simpler, shallower, and perhaps evolutionarily older mechanism for local, word-by-word predictions. Indeed, the co-existence of two distinct mechanisms was already identified in the case of the processing of sequences of non-linguistic items (23–29). Specifically, using the Local-Global Paradigm, a variant of the oddball auditory paradigm, Bekinschtein et al. (23) showed that transition-based (local) processing can be distinguished from that of chunking (global) and from possibly higher levels, each eliciting distinct violation signals in different brain areas and timings (30–35).

The co-existence of two such distinct mechanisms was also identified in artificial neural language models. Lakretz et al. (36, 37) showed that, during training, neural language models develop two types of mechanisms to process word dependencies in language. One mechanism was shown to be sensitive to the latent hierarchical syntactic structure of sentences, and therefore capable of computing long-distance dependencies, whereas the other mechanism was only sensitive to local sequential transitions between words and therefore structure agnostic.

Finally, there is also a long history in psycholinguistics already pointing to the existence of two types of cognitive mechanisms involved in the processing of long-range agreements. Much behavioral work from classic psycholinguistics has shown that humans succumb, at least in part, to local effects that interfere with higher, hierarchical, processing of sentences. In a classic work, Bock and Miller (38) showed that human participants make many long-range agreement errors on sentences such as (1)

(1) The **boy** near the *girls* **likes** climbing

*Det N*_1_ *P Det N*_2_ *V N*

In this example, the main (‘boy’) and embedded (‘girls’) nouns have different numbers (one is singular, the other plural). Such incongruent sentences were found to be more error-prone than when the two nouns are congruent (e.g., both singular). This local behavioral interference effect was later replicated in many studies across a variety of syntactic structures (e.g., 39–43).

Such a behavioral effect could, however, arise from various underlying reasons. In particular, two possible explanations for local interference are discernible: Let *N*_1_, *N*_2_, and *V* denote the main noun (e.g., ‘boy’), embedded noun (‘girls’) and verb (‘likes’) in sentences of the form 1 (such that *V* should agree with *N*_1_). One explanation for the local interference of noun *N*_2_ could be a *memory interference* between *N*_1_ and *N*_2_: the mental representations of the two nouns would interact during sentence processing, such that the memorized number of *N*_1_ would occasionally be overridden by the more recent number of *N*_2_, in memory encoding or retrieval. Such noun-noun interference would be in line with theories from psycholinguistics, such as the cue-based retrieval theory (40). A second alternative explanation for local interference could attribute it to *conflicting prediction signals* from *N*_1_ and *N*_2_ onto *V* . The correct, long-distance, or global expectation for the corresponding grammatical number of *V*, based on *N*_1_′ *s* number, would be affected by a recent expectation generated by the more recent *N*_2_. Such a local influence would be in line with findings from non-linguistic auditory stimuli (the local effect observed by Bekinschtein et al. (23)) as well as from the above-described neural language models, where two neural predictive mechanisms coexist and can, in some cases, generate opposing, competing predictions.

Since behavioral evidence alone is compatible with both explanations, here, we tested neurally the possible coexistence of two neural mechanisms in the human brain, one which is sensitive to the latent structure of sentences, and another, simpler, possibly evolutionarily older, which would be sensitive to mere word transitions. If the predictions from the models and from the processing of auditory stimuli hold in the case of language processing in humans, then we would expect to find neural evidence for the two distinct types of mechanisms. Alternatively, it is possible that language processing is entirely based on hierarchical mechanisms and that behavioral interference can be solely accounted for by interference between the two representations of *N*_1_ and *N*_2_ in memory.

To test these hypotheses, we created a 2×2 design in which both local transitions and long-distance relations of agreement were manipulated orthogonally and could be independently respected or violated. We recorded neural activity from both human participants (*n* = 22, magnetoencephalography; MEG) and a neural language model and studied whether the neural signatures of the two levels can be disentangled. In both cases, we used temporally resolved multivariate decoding techniques (32) to probe the existence of long-distance, local, and memory-interference mechanisms.

To anticipate the results, in the model, we found evidence for two distinct neural effects, which correspond to the two co-existence of long-distance and local mechanisms, corroborating our previous results (36, 37). In human brain signals, however, only a main effect related to the hierarchical structure of sentences was found. While this effect was modulated by noun-noun congruity, there was no evidence of any brain signal reflecting a local sequential effect. These findings suggest that, unlike low-level processing of auditory stimuli, in sentence processing, once words enter the language system, computations are dominated by structure-based processing and are largely robust to sequential effects. Furthermore, they suggest that memory interference between representations of incongruent nouns is the dominant factor underlying local interference observed behaviorally in psycholinguistic work.

## Methods

A total of 22 participants with normal or corrected to normal vision were included in the M/EEG experiment. The number of the selected participants was based on previous studies in subject-verb agreement experiments (44), where an average of 23.3 participants was shown to be sufficient to detect subject-verb agreement violations, with a few studies reporting results with less than 15 participants. According to Molinaro et al. (45), a sufficient amount of participants for an agreement study is of 20. In compliance with the institutional guidelines, all participants gave written, informed consent prior to the experiment and were compensated with 100 Euros for their participation. The participants were native English speakers and prior to the participation, the subjects had to perform an online sentence reading task (Dialang^1^ reading & structures task—subjects accepted with a placement greater than C1 in both tests). The procedure and the consent were approved by the local ethical committee.

### Stimuli

To contrast hierarchical and sequential mechanisms, we make use of long-range grammatical agreements as in sentence 1, where we manipulate:

1. The *structural* relation between *N*_1_ and *V*.
2. The *sequential* intervention of *N*_2_ on the processing of *V*.

These two dimensions span the two-by-two design of the paradigm, and its corresponding two main effects are defined as follows (Figure 1): A *structural effect*, which contrasts conditions in which *N*_1_ and *V* agree and disagree on grammatical number. In the case of disagreement, a syntactic violation occurs, and neural response to this violation is expected, as was extensively studied in past studies (46, 47). The second one is a *transition effect*, which contrasts conditions in which *N*_2_ and *V* match and mismatch with respect to grammatical number. The transition effect corresponds to local word transitions and was not identified in neural recordings thus far. Following results from the classic local-global paradigm and from simulations in neural language models, we hypothesized that a local number mismatch would violate local word-transition expectations, which, in turn, would generate an identifiable neural response, independently of whether a syntactic violation simultaneously occurs. For example, in sentence 1, the frequency of ‘girls likes’ is two orders of magnitude smaller than that of ‘girl likes’ (logfrequency = -8 and -6, respectively; Google’s n-gram). Lowlevel brain regions of the language network might be sensitive to such transition probabilities, in which case, a greater neural response is predicted for the low -compared to the high-frequency word pair. This is consistent with a predictive-coding framework (48–50), which suggests that cortical circuits form an internal model of input sequences and that this model continuously generates predictions about upcoming items, confronting them with incoming stimuli. The local effect would thus reflect a prediction error that results from an internal model based on transition probabilities.

**Fig. 1.**
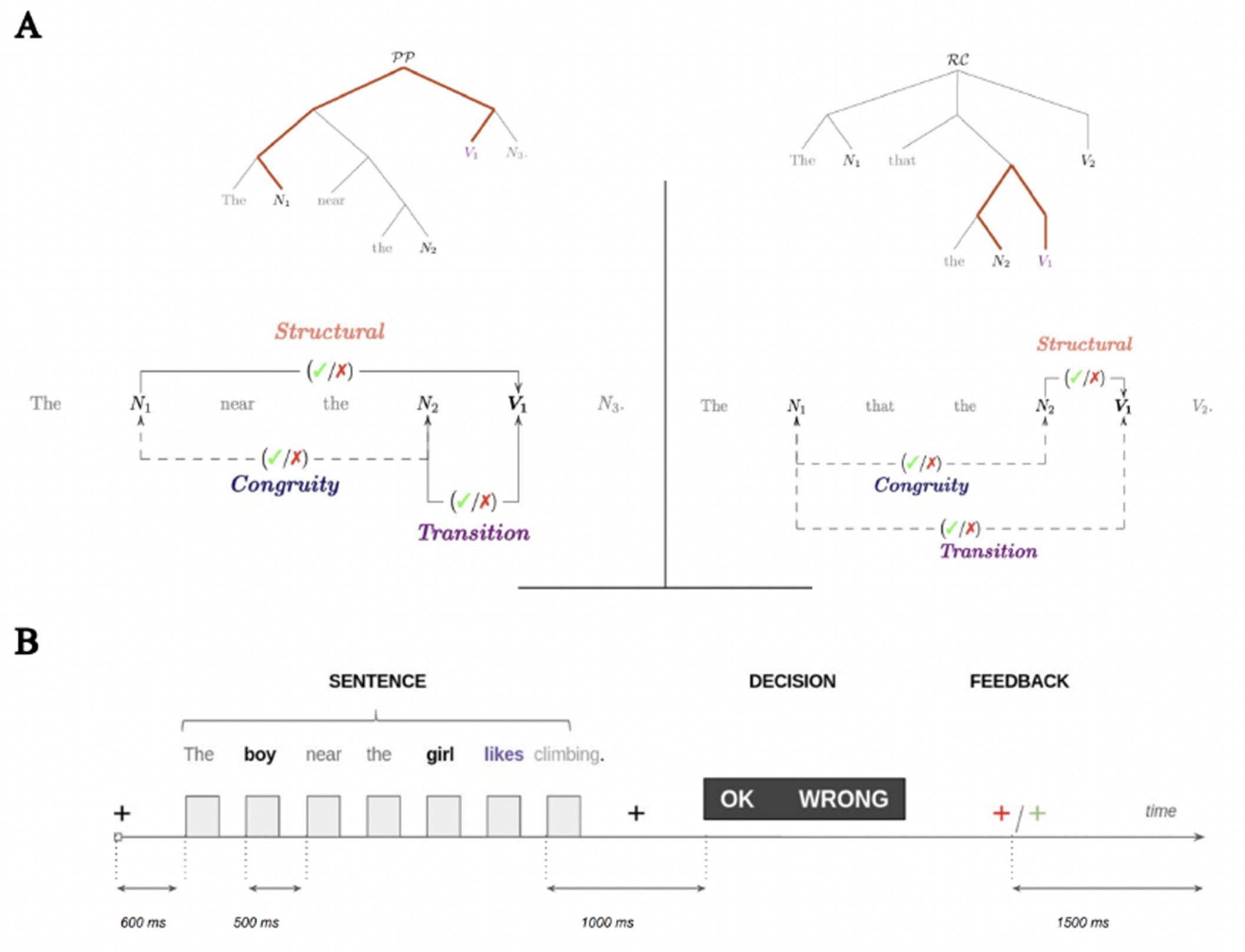
Structural vs. linear intervention in sentence processing—experimental design and paradigm. To disentangle two possible types of processing during sentence comprehension, the experimental design contrast: (i) a structural dependency between a target verb and a noun, which either holds, or creates a violation at the verb, and (ii) a linear (sequential) interaction between the target verb and another noun, which either facilitates or interferes with verb processing. (A) Tree representations of the two sentence constructions explored in the experiments. Below, is an illustration of the main effects of the design: Structural effect (orange), which depends on the syntactic relation between the main subject and target verb (colored path in the tree representation). Transition effect (magenta), which refers to the (mis)match between the target verb and a linearly intervening noun (attractor), with respect to either grammatical number or animacy. congruity effect, which refers to the (mis)match between the two nouns; In the left construction, the structural effect is long-range and the transition (linear) one is short-range. On the right construction, it is the opposite. (B) Experimental Paradigm: subjects were presented with sentences in a rapid serial visual presentation (RSVP), and their task was to report whether the sentences are grammatically correct. At the end of each trial, visual feedback on their performance was given.

**Fig. 2.**
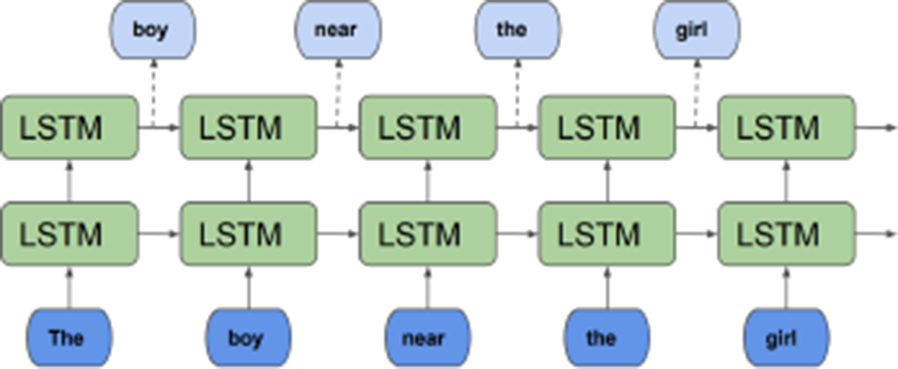
Graphical description of a two-layer recurrent neural language model with LSTM cells (not discussed here; see, e.g., 61). At each time step, the model processes an input word and outputs a probability distribution over the potential next words in the sentence. The prediction of the output word depends on both the input word and on the previous state of the model, which serves as longer-term context (the horizontal arrows in the figure represent the *recurrent* connections carrying the previous state through).

In the construction with a PP, the global (structural) and local (transition-based) effects are, however, correlated with linear proximity. That is, the global effect between the subject *N*_1_ and verb *V* is long-range, whereas the local effect between the intervening noun *N*_2_ and verb is short-range. To decouple syntactic dependency and linear proximity, we included a control construction in the design, in which syntactic dependency and linear proximity are reversed by the replacement of only a single word. Specifically, we replaced the PP with an object-relative clause (ObjRC; Figure 1, Table 1):

**Table 1.** Design and prototypical examples. The experiment utilizes two linguistic constructions, a Prepositional Phrase (PP) and an Object Relative Clause (ObjRC) as well as two features of interest (Number & Animacy). The manipulation of the violation factor corresponds to the structural effect.

| Construction | Template | Examples | Violation | Congruity |
| --- | --- | --- | --- | --- |
| *PP – Number* | Det $N_1$ P det $N_2$ $V_1$ $N_3$ | The boy near the girl likes climbing. | No | Yes |
|  |  | The boy near the girls likes climbing. | No | No |
|  |  | The boy near the girls like climbing. | Yes | No |
|  |  | The boy near the girl like climbing. | Yes | Yes |
| *ObjRC – Number* | Det $N_1$ Adv det $N_2$ $V_1$ $V_2$ | The boy that the girl likes leaves. | No | Yes |
|  |  | The boy that the girls like leaves. | No | No |
|  |  | The boy that the girls likes leaves. | Yes | No |
|  |  | The boy that the girl like leaves. | Yes | Yes |
| *PP – Animacy* | Det $N_1$ P det $N_2$ $V_1$ $N_3$ | The boy near the girl likes climbing. | No | Yes |
|  |  | The boy near the car likes climbing. | No | No |
|  |  | The boy near the car rusts badly. | Yes | No |
|  |  | The boy near the girl rusts badly. | Yes | Yes |

**Table 2.** The exact stimuli presented in the first run of the experiment for the first subject of the study.

| sentence | violIndex | condition_type |
| --- | --- | --- |
| The vet that the author knows dies. | 0 | syntactic |
| The teachers that the dentist loves sneeze. | 0 | syntactic |
| The waiter that the doctors dislikes turns. | 1 | syntactic |
| The judges by the doctors likes painting. | 1 | syntactic |
| The tractors near the actors dislike farming. | 1 | semantic |
| The lawyers by the doctor hate skiing. | 0 | syntactic |
| The taxi by the teacher dislikes fencing. | 1 | semantic |
| The nurse by the vets hates walking. | 0 | syntactic |
| The judge by the vets hates knitting. | 0 | syntactic |
| The painter that the teacher dislike moves. | 1 | syntactic |
| The vets near the lawyers hate driving. | 0 | syntactic |
| The bankers near the nurses hates shopping. | 1 | syntactic |
| The doctors near the doctor loves fishing. | 1 | syntactic |
| The athletes beside the jeeps love boxing. | 0 | semantic |
| The doctor by the judge honks loudly. | 1 | semantic |
| The cars by the lawyers love shopping. | 1 | semantic |
| The tailors that the barber love swim. | 1 | syntactic |
| The plumber beside the nurse loves driving. | 0 | syntactic |
| The actor that the painters likes coughs. | 1 | syntactic |
| The van near the truck knows Japanese. | 1 | semantic |
| The scooter beside the crane brakes abruptly. | 0 | semantic |
| The lawyer that the doctors dislike cries. | 0 | syntactic |
| The author near the nurse know Russian. | 1 | syntactic |
| The car near the actor hates fishing. | 1 | semantic |
| The jeep beside the doctor brakes noisily. | 0 | semantic |
| The lawyers that the baker admire cry. | 1 | syntactic |
| The athletes that the painter hates sit. | 0 | syntactic |
| The bakers by the bankers like spinning. | 0 | syntactic |
| The teacher near the truck hates cooking. | 0 | semantic |
| The jeeps by the buses hate cooking. | 1 | semantic |
| The taxis beside the trucks admire gratitude. | 1 | semantic |
| The actor that the doctor loves smiles. | 0 | syntactic |
| The athlete that the teacher dislike sneezes. | 1 | syntactic |
| The pilot beside the lawyer likes acting. | 0 | syntactic |
| The bankers that the painters loves die. | 1 | syntactic |
| The authors that the nurses likes die. | 1 | syntactic |
| The farmers by the banker like farming. | 0 | syntactic |
| The truck near the bus skids suddenly. | 0 | semantic |
| The actor beside the actors dislike driving. | 1 | syntactic |
| The tailor that the judges like prays. | 0 | syntactic |
| The actor beside the dentists like cycling. | 1 | syntactic |
| The scooters beside the taxis skid loudly. | 0 | semantic |
| The tailors that the doctors love leave. | 0 | syntactic |
| The actors that the lawyers like leave. | 0 | syntactic |
| The vans beside the buses skid abruptly. | 0 | semantic |
| The doctors by the cranes dislike spinning. | 0 | semantic |
| The teachers near the baker knows Arabic. | 1 | syntactic |
| The plumber beside the painter know French. | 1 | syntactic |

(2)**ObjRC**: “The boy that the **girls like**… “(“*Det N*_1_ *that det N*_2_ *V*”),

In this case, the structural dependency (in bold) is now between *N*_2_ and *V*, whereas the intervening noun is the distant *N*_1_. The difference between the two sentences 1 and is minimal — only at the third word (‘near’/’that’), and the number of words that precede the target verb *V* is the same. This allows us to test the impact of the length of the subject-verb dependency on the global effect, and the impact of the proximity between *N*_2_ and *V* on the local effect. In particular, to test the prediction that a neural response to local-transition violations would occur only in the case of a PP but not in the ObjRC case.

So far, the examples shown contained variations of a sentence with respect to the feature of grammatical number. The number feature and the corresponding agreement phenomena are generally perceived as a proxy for syntactic processing. However, violation responses are known to vary depending on whether the violation is semantic or syntactic. While syntactic violations typically generate a late positive neural response P600 (51), semantic violations were found to elicit an earlier negative response N400 (52). We therefore further manipulated the type of feature in the agreements, and it includes sentences of the form 1, in which violations are with respect to animacy, for example:

(3)**PP**: “The **boy** near the car **likes** climbing” (“*Det N*_1_ *near det N*_2_ *V* “),

In these sentences, the subject *N*_1_ and verb *V* always agree on the number, however, the sentence contains a semantic violation, which is either local (‘car likes’), or global (as in, “The boy near the girl rusts badly”).

Table 1 summarizes the three constructions of the design: PP-Number, PP-Animacy, ObjRC-Number, along with example sentences. Note that due to a time constraint on the entire experimental duration, we did not include a fourth case in the design, which includes sentences with an object-relative clause and semantic violations. For each of the three constructions, we generated 16 stimuli per block for a total of 10 blocks. Half of these stimuli contained a violation. The stimuli were generated using a simple algorithm that sampled without replacement words from the lexicon. Each participant was presented with a different set of stimuli. The lexicon consisted of 19 animate nouns, 7 inanimate nouns, and a total of 15 verbs. The stimuli were controlled for low level features such as length and unigram frequency. ^2^ Each participant was presented with an equal number of sentences, ensuring that the count of sentences with a singular first noun matched those with a plural first noun. Similarly, for the semantic conditions, the number of sentences featuring an animate first noun was identical to those with an inanimate first noun. Additionally, a sentence could not have the same noun in both singular and plural form (e.g: a sentence such as: “The boy near the boys…” was forbidden). More examples of stimuli are presented in appendix C.

### Experimental Paradigm

The participants undertook a rapid serial visual presentation (RSVP) reading task and were asked to report whether a sentence contained a grammatical or semantic violation by pressing a button on a MEG response device.^3^ To verify that participants understood the task, prior to recording, they went through a short training phase (10 minutes). The task was divided into 10 equal runs, where each run contained the same number of trials (*n* = 48). A 600*ms* fixation cross interval preceded the onset of the first word (Figure 1B). All the sentences had the same length. The words were presented with a stimulus onset asynchrony (SOA) of 500*ms*. After the offset of the last word and following a time interval of 1s, a decision panel with the words “OK” and “WRONG” appeared on the screen. To control for motor preparation, the location of the words (left or right) was randomized at each trial. As soon as the participants stated their decision, the decision panel disappeared and the subjects received immediate visual feedback on their performance. If their response was correct, they were presented with a green cross, otherwise with a red one. Decision duration was limited to 1.5*s*, after which a blue fixation cross appeared and the experiment continued. The interval to the next trial (ITI) was 1.5*s*. All time intervals were set to multiples of the video projector refresh rate (60*Hz*).

### M/EEG recordings

Recordings took place in two different MEG centers: NeuroSpin in Saclay, France (*N* = 15), and ICM in Paris (*N* = 7). Participants performed the task while sitting in an electromagnetically shielded room. Brain magnetic fields were recorded with a 306-channel, whole-head MEG by Elekta Neuromag® (Helsinki, Finland), in 102 triplets: one magnetometer and two orthogonal planar gradiometers. In NeuroSpin and ICM, EEG was recorded with a 60 and 64-channel MEG-compatible Neuromag EEG cap, respectively. The brain signals were acquired at a sampling rate of 1000*Hz* with a hardware highpass filter at 0.03*Hz*. Eye movements and heartbeats were monitored with vertical and horizontal electrooculogram (EOGs) and electrocardiograms (ECGs). The subjects’ head position inside the helmet was measured at the beginning of each run with an isotrack Polhemus Inc. system from the location of four coils placed over frontal and mastoïdian skull areas. All EEG sensors were digitized as well.

### Preprocessing

Bad sensors per sensor type were automatically detected at the run level based on a variance criterion. Channels in which the variance exceeded the median channel variance by 6 times, or was less than the median variance divided by 6, were marked as bad. A visual inspection was followed to verify the detection accuracy. Prior to the variance detection, oculomotor and cardiac artifacts were removed at the run level, using signal-space projection (SSP) implemented with MNE Python (53, 54). To compensate for head movement and reduce non-biological noise, the MEG data were Maxwell-filtered (55) using the implementation of Maxwell filtering in MNE Python. The bad EEG sensors were interpolated using the spherical spline method (56) implemented in the same package. Following Maxwell filtering, the linear component of the data was removed, and the time series were clipped at the upper and lower bound values of (-3,3) interquartile range (IQR) around the median. The data were then bandpass filtered between 0.4 and 50 Hz using a linear-phase FIR filter (hamming) with delay compensation, implemented in MNE-python version 0.16 (53). Finally, the continuous time series were segmented into 3.5s epochs of interest (first word onset to panel onset) and the SSP procedure was applied to the epoched data to remove heart-beats and ocular motions.

### Decoding Analyses

We used a temporally-resolved decoding approach to classify neural activity from two conditions at the trial level (32, 57). These analyses were implemented in MNE-python version 0.16 (53). Prior to model fitting, the data were standardized using the Scikit-Learn package (58). We used a linear classifier (logistic regression) with default Scikit-Learn parameters. The evaluation metric was the Area Under the Curve (AUC). The estimator was trained and tested on data from the same condition. To prevent overfitting, we used a 5-fold stratified cross-validation procedure.

### Statistical Analyses

The reported statistics for the behavioral data are performed with a mixed-effects logistic regression with the subject number as a random variable. These were performed using the lme4 package in R (59). The reported statistics for the neural data correspond to group-level analyses and were performed using the Statsmodels package in Python3 and in MNE-python version 0.16 (53). The statistical significance of the decoding performance over time was evaluated and corrected for multiple comparisons using a cluster-based permutation approach (60), using a total of 1000 permutations. The significance threshold (alpha level) for all analyses was set to 0.05.

### Neural Language Models

In the computational experiments, we use a Neural Language Model (NLM). An NLM is a language model implemented by a neural network, defining a probability distribution over sequences of words. It factorizes the probability of a sentence into a multiplication of the conditional probabilities of all words in the sentence, given the words that precede them:

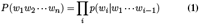

This type of language model can thus be used as *next-word predictor*: given the preamble of a sentence, it outputs a probability distribution over potential next words. We exploit this fact in our experiments.

### Model description

The specific NLM we use in our experiments is the English NLM made available by Gulordava et al. (62)^4^. It is a recurrent LSTM language model (63), consisting of two layers, each with 650 Long-Short Term Memory units (64), input, and output embedding layers of 650 units and input and output layers of size 50000 (the size of the vocabulary). The weights of the input and output embedding layers are not shared (65). The last layer of the model is a softmax layer, whose activations sum up to 1 and as such corresponds to a probability distribution over all words in the NLM’s vocabulary.

### Model training

The weights of an NLM are typically tuned by presenting them with large amounts of data (a *training corpus*) and providing them feedback on how well they can predict each next word in the running text. This allows them to adjust their parameters to maximize the probabilities of the sentences in the corpus. Our NLM was trained on a sample of the English Wikipedia text, containing 100M word tokens and 50K word types. Further details can be found in Gulordava et al. (62).

## Results

We first present the behavioral results from the experiment, based on the performance of the subjects in a forced-choice, violation-detection task (Figure 1C). We then present classification results in a time-resolved manner on the main effects of the design, for both humans (MEG & EEG) and artificial language models.

## Behavioral Results

Figure 3 shows the mean error rates across all participants for the three main constructions. We present the error rates with respect to the structural and congruity effects, whereby congruity effect we refer to whether the two nouns *N*_1_ and *N*_2_ agree on number or not. In the following, we report results from a mixed-effects logistic regression model (see Methods).

**Fig. 3.**
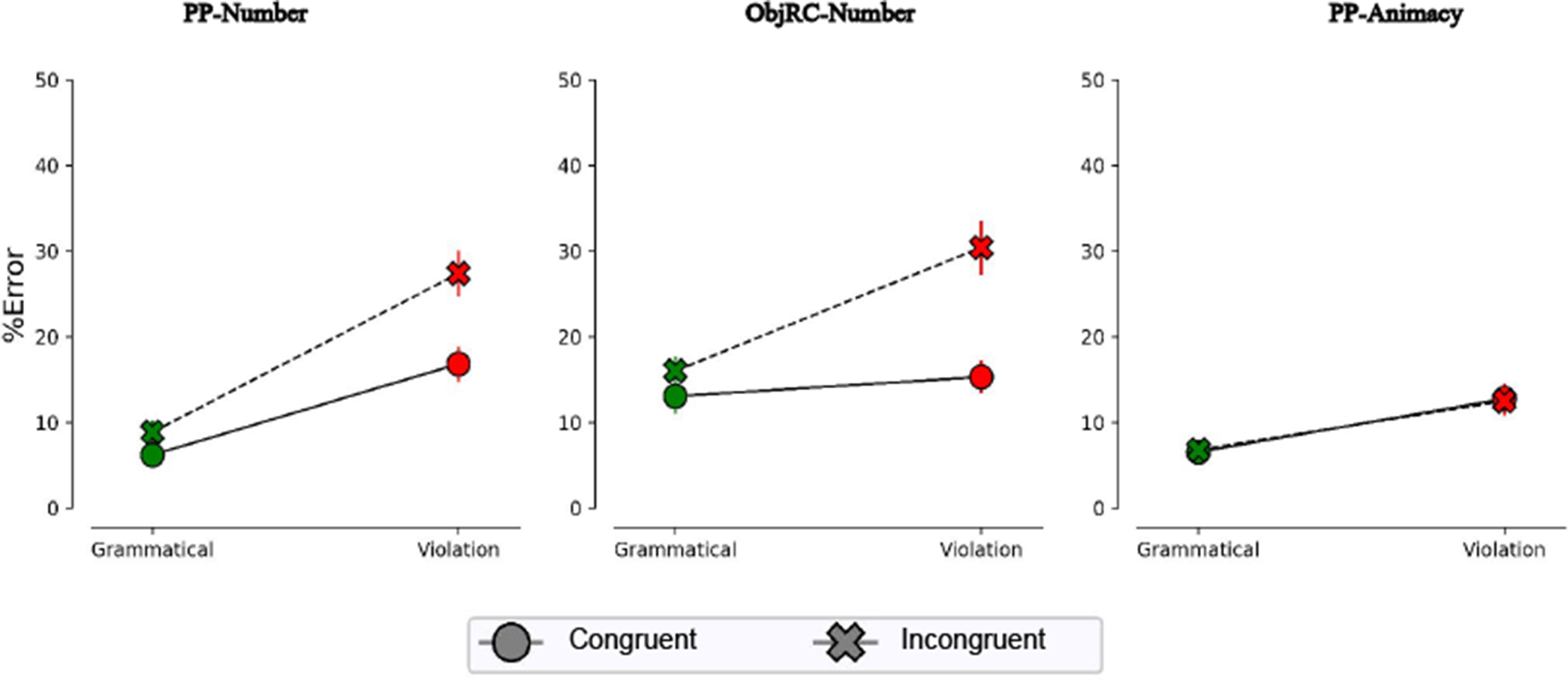
Behavioral results. Interaction plots for the Structural and congruity effects (N=22). The main structural and congruity effects, as well as their interaction, are significant in the number of constructions (*p <* 0.05). In the animacy condition, only the structural effect is significant. The error bars indicate the standard error of the mean (SEM) calculated across participants.

We first tested whether the *structural effect* (Figure 1C) is present at the behavioral level. Indeed, this effect was significant across all constructions. This indicates that participants made more errors in detecting a violation, compared to affirming that a sentence was grammatical, which is akin to ‘grammatical illusion’ (66). The effect was stronger for PP-Number (*b* = 1.8488, *SE* = 0.1659, *z* = 11.141, *p < .*001), followed by ObjRC-Number (*b* = 1.1332, *SE* = 0.1309, *z* = 8.659, *p < .*001), and finally PP-Animacy (*b* = 0.74834, *SE* = 0.17491, *z* = 4.278, *p < .*001).

We then examined the effect of *congruity* on error rate. Notably, for PP-Number the effect was significant (*b* =−1.0592, *SE* = 0.2268, *z* =−4.671, *p < .*001). For ObjRC-Number, the congruity effect was also significant (*b* =−1.1894, *SE* = 0.1885, *z* = −6.308, *p < .*001). However, for PP-Animacy, the congruity effect was not significant (*b* =−0.03540, *SE* = 0.24583, *z* =−0.144).

Finally, we investigated the *transition effect*, which corresponds to a mismatch between the intervening noun and the target verb, e.g., *N*_1_ and *V* in (2). Notably, for PP-Number this effect was significant (*b* = 0.3935, *SE* = 0.1905, *z* = 2.066, *p* = .0388). For ObjRC-Number, the effect was marginally significant (*b* = 0.2719, *SE* = 0.1416, *z* = 1.921, *p* = .0548). However, for PP-Animacy, the transition effect was not significant (*b* = 0.03901, *SE* = 0.19601, *z* = 0.199). The transition effect for number is consistent with ‘grammatical asymmetry’ (43), where participants make more agreement errors on ungrammatical compared to grammatical sentences.

In summary, at the behavioral level, we observed a significant structural effect, for both constructions (PP & ObjRC) and both features (grammatical number and animacy). Transition and congruity effects were significant for both constructions but for grammatical number only. And, finally, participants made more errors in agreement in an embedded clause compared to the prepositional phrase.

### Structural but not Transition Effects are Decodable in Human Data

The transition and congruity effects for grammatical number in the behavioral data suggest that the representation of the two nouns, *N*_1_ and *N*_2_, interact during sentence processing. There could be two types of interactions: Interference in memory. An incongruent number of *N*_2_ interferes with that of *N*_1_ in memory; (2) Conflicting prediction signals. A (global) expectation for the corresponding grammatical number of *V*, based on 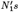 number, is affected by a (local) expectation generated by *N*_2_. However, the behavioral results cannot tell apart the two, since both alternatives predict the same behavioral outcome. Therefore, we next tested whether these effects are traceable in the neural data.

We sought the effects in both the model and in humans. We presented the same stimuli to human participants and to the model, and recorded network activity after the presentation of each word. For humans, neural activity was recorded with a magnetoencephalography (MEG) machine. For the model, we extracted the hidden activity of all recurrent units of the network (Methods).

To identify the main effects in the data, we used standard decoding techniques: for each effect, at each time point, a linear binary classifier was trained to separate trials from the two conditions, and then tested on unseen data in a cross-validation manner. Figure 4 shows the decodability of the main effects for both the artificial (panel A) and human data (panel B).

**Fig. 4.**
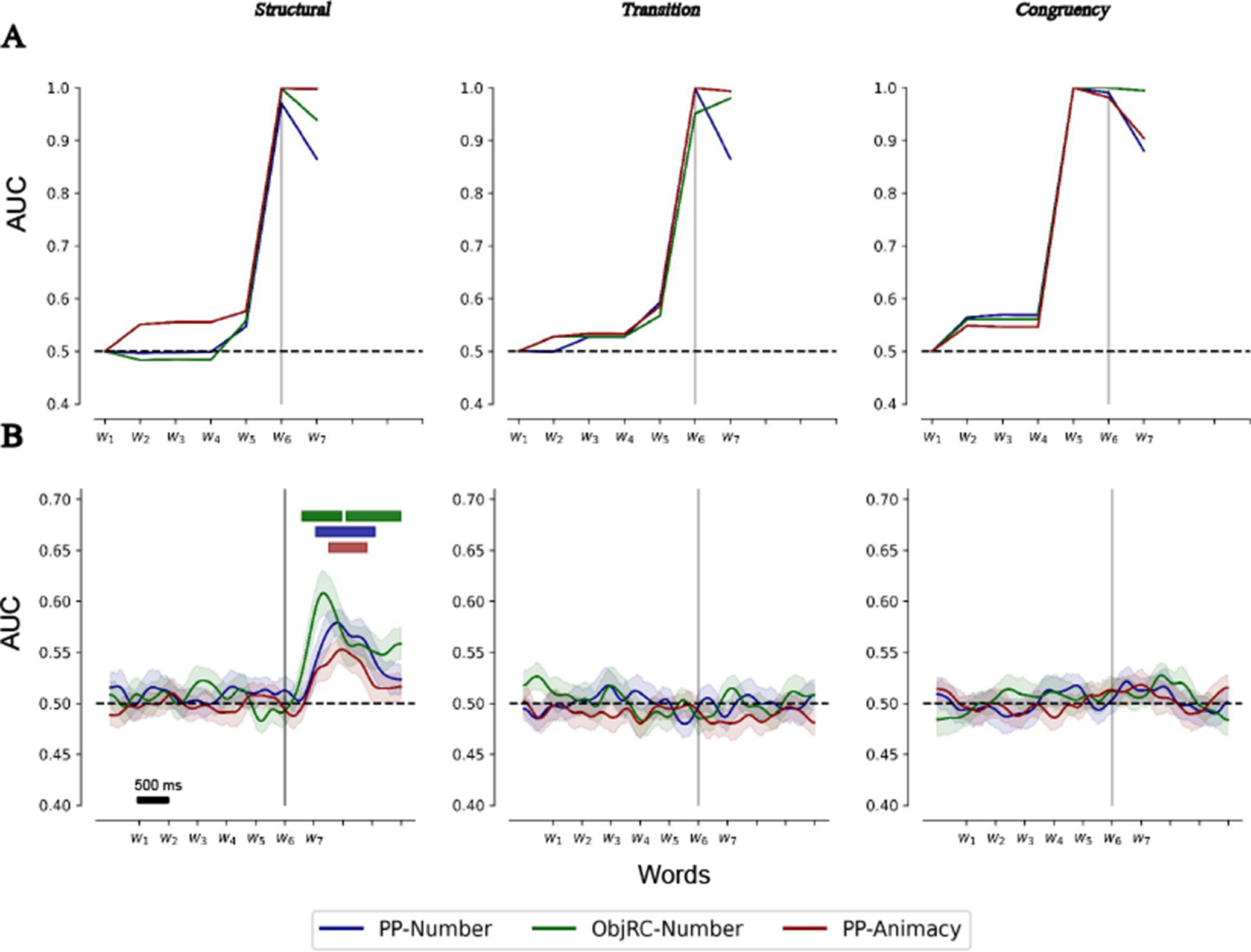
Structural but not linear effects are decodable in human data. (A) Decoding of the main effects originating from the activations of an LSTM architecture. All main effects are decodable. (B) Neural decoding of the three main effects using all sensor types (magnetometers, gradiometers, and EEG). A different decoder was trained on each time point and construction, and evaluated using the Area Under the Curve (AUC). Significant decoding performance was found for the structural effect only. Color stripes indicate statistically significant time intervals (*p <* 0.05; corrected—spatio-temporal clustering permutation test). The decoding for the main effects of transition and congruity remained at the chance level until the end of the time window of interest. Results are shown for correct responses only, and the data was smoothed with a 100ms moving Gaussian kernel for visualization purposes only.

For the model (Figure 4A), for all three constructions, all effects were decodable with high performance, measured in terms of the Area Under of Curve (AUC). The structural effect reached full decodability after the onset of the target word. Indeed, prior to the onset of the target verb, the model cannot predict the grammaticality of the sentence. The transition effect reached full decodability also after the onset of the target word. Here too, prior to verb onset, a mismatch between the verb and the non-head noun cannot be predicted. Finally, the congruity effect was decodable already after the onset of the second noun. Indeed, information about feature mismatch between the two nouns is already available at this time point.

For the MEG data, and in contrast to the model, only the structural effect was decodable. The onset and the peak decodability of the effect varied across constructions. The effect becomes significant first, for the ObjRC-number (*t* = 300*ms*), then for the PP-Number (530*ms*), and lastly for the PP-animacy (760*ms*). The significance of the decodability was calculated based on cluster-based permutation testing (60; Methods). The decodability of the structural effect reached its highest value for the ObjRC-number construction (*AUC* : 0.61), followed by the PP-Number (*AUC* : 0.58) and finally by PP-animacy (*AUC* : 0.55).

We observed a discrepancy between the behavioral results, the decoding from the neural-network model, and the Decoding from the neural data At the behavioral level, the structural effect was the dominant factor, and effects emerging from the interaction withs the intervening noun were significant only for the number feature. In the model, we observed the same sensitivity across all three main factors, namely, significant effects arising from the intervening noun for both the animacy and the number feature. Lastly, in the human neural data, only the structural effect was decodable, whereas the performance of decoding the other effects remained at chance level.

In summary, we found strong positive evidence from the neural data for a structural effect in humans, but negative evidence for the transition effect. This goes against the conflicting-prediction-signals hypothesis and is in favor of the memory-interference one. We therefore next sought to identify positive evidence in favor of the memory-rinterference hypothesis.

### The influence of the attractor on the structural effect

In our search for positive evidence for the noun-noun congruency effect, we reasoned that this effect should modulate the long-distance congruity effect: on trials where the two nouns are incongruent, the memory of the first noun *N*_1_ would suffer and therefore the brain’s response to a long-distance violation would be weaker and/or delayed compared to trials with two congruent nouns. To test this idea, for each construction, we first trained a linear binary classifier on the structural effect, and then, separately, tested its predictions on different conditions in the test data (Figure 5A). At training time, the classifier was trained to separate all violation trials from non-violation trials, regardless of whether they were congruent or not. Then, at test time, the classifier was separately tested on unseen trials that were either congruent (continuous lines) or incongruent (dashed).

**Fig. 5.**
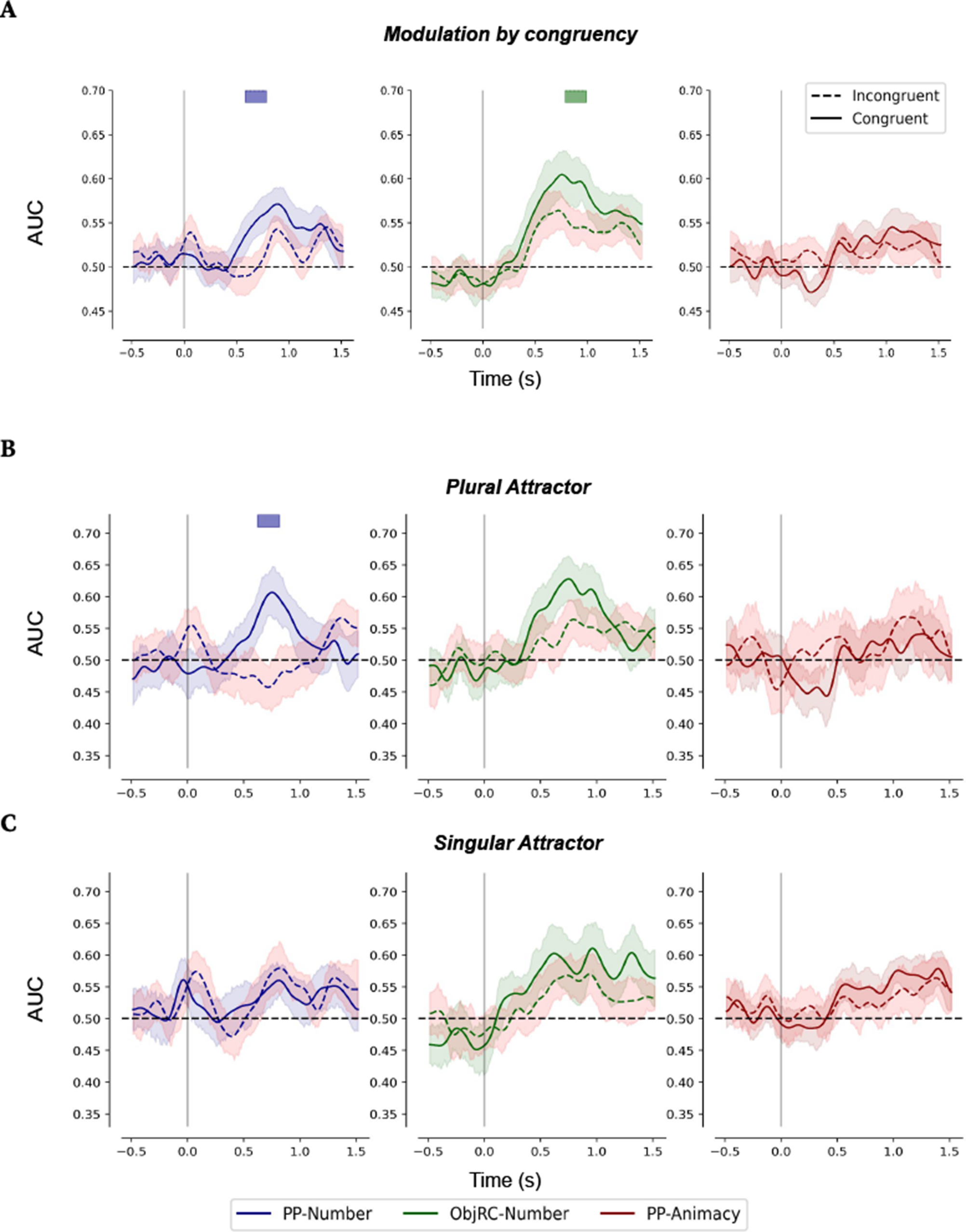
Modulation of the structural effect by congruity and neural correlates of the markedness effect. (A) A classifier was trained on the structural effect (grammatical vs. ungrammatical trials), per construction, and subsequently evaluated on the structural effect when contrasting the congruent (continuous lines) and incongruent (dashed) trials. Color stripes indicate statistically significant time intervals (*p <* 0.05; corrected using spatio-temporal clustering permutation test). Congruity was found to modulate the structural effect for both PP-number and ObjRC-number, but not for PP-Animacy. (B&C) For each construction, a classifier was trained on the structural effect, and subsequently tested on the same effect when trials were split for both congruity and attractor number. For PP-number, a statistically significant difference was found for the plural but not the singular case (*p <* 0.05; corrected—cluster-based permutation test). For ObjRC-Number and PP-Animacy, no significant differences were found. Results are shown for correct responses only.

**Fig. 6.**
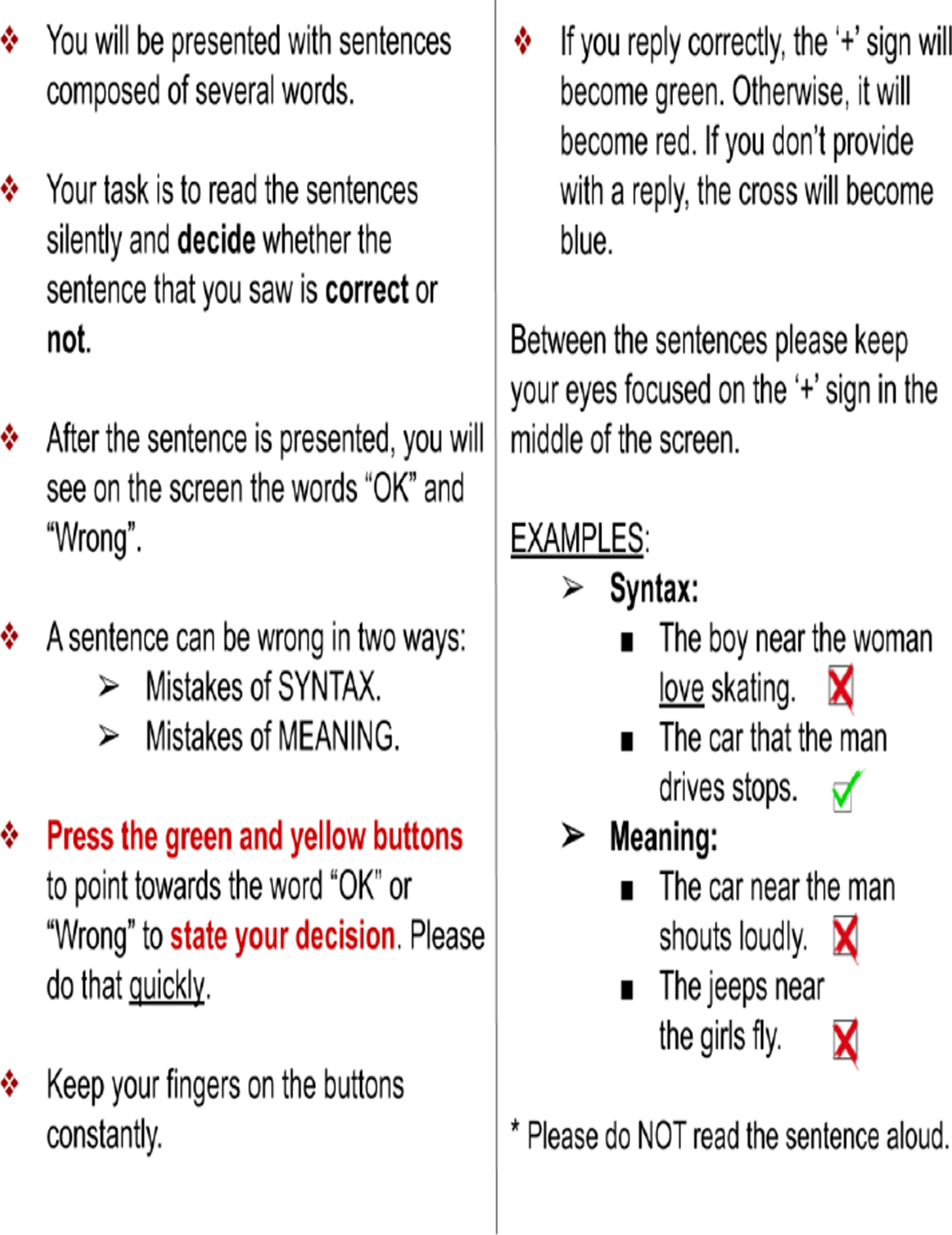
The exact instructions as they were presented to the subjects prior to the beginning of the experiment.

**Fig. 7.**
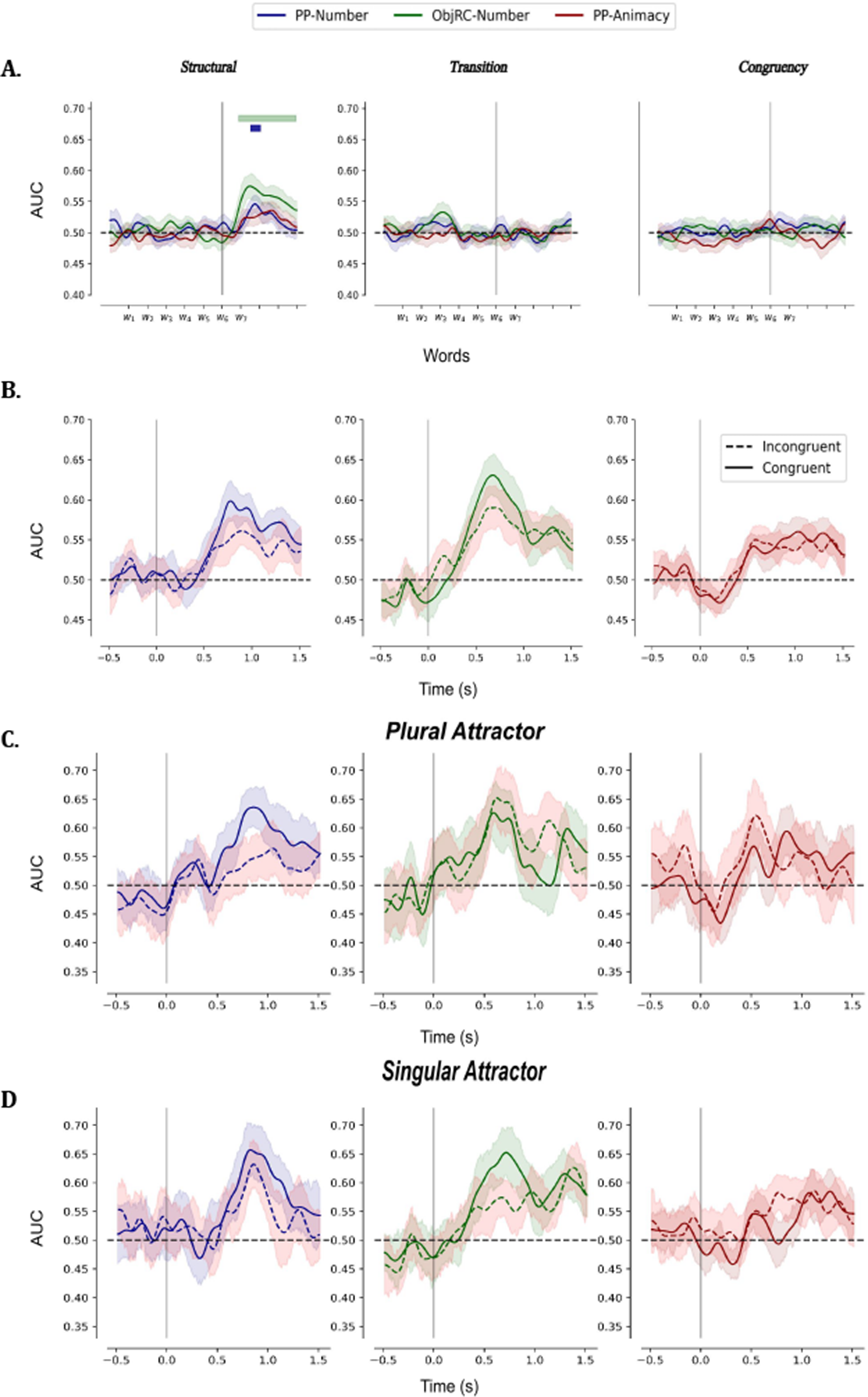
A. Structural but not linear effects are decodable in human data (all responses; see Figure 4). (B-D) Modulation of the structural effect by congruity and neural correlates of the markedness effect (correct responses only; See Figure 5).

For PP-Number, we found that the structural effect was modulated by congruity. The difference between the congruent and incongruent trials became significant at 580*ms* after the onset of the target and was sustained for 200*ms*. For ObjRC-number, the structural effect was also modulated by congruity. This modulation became significant later compared to the one for the PP-Number construction, starting at 780*ms* and up until 980*ms* after the target onset. Finally, for PP-animacy, the structural effect was not modulated by congruity, in complete agreement with the behavioral results, where no behavioral interference was observed.

In sum, these results provide positive evidence for the congruency effect and in favor of the memory-interference explanation, where behavioral interference arises from interference between the two noun representations in memory.

### The neural correlates of the markedness effect

Our design allowed us to study also a well-established phenomenon related to the processing of long-range agreement, known as the *markedness effect*. In classic work on grammatical agreement, it was observed that for sentences with a long-range subject-verb agreement, participants make more errors if the attractor is plural compared to singular (38). For example, comparing the two following sentences,

1. “The boy near the girls “.
2. “The boys near the girl “.

On average, participants made more errors on (1) compared to (2). This phenomenon is known as the markedness effect, since in English, plural is the marked form (English: Bock and Miller 38, Wagers et al. 43, Eberhard 67; Italian: Vigliocco et al. 68 Spanish: Bock et al. 69, Lago et al. 70 French: Franck et al. 71 Russian: Lorimor et al. 72).

Thus far, this phenomenon has beenthrough behavioral results We thus took advantage of our data to investigate whether we could identify a neural correlate of this phenomenon.

We previously saw that the congruity effect modulated the structural effect (Figure 5A). To investigate the markedness effect, we took this analysis one step further: as before, we trained the classifier on the structural effect, but now we analyzed its performance separately for each value of congruity and attractor number (Figure 5B&C).

When examining the modulation of the structural effect by congruity, we observed no difference between the congruent and incongruent trials in the case of a singular attractor, for none of the three constructions. However, when examining the same effect for the plural attractor, the decodability of the congruent trials was significantly higher compared to the incongruent ones. This difference was significant for the time interval between 620*ms* and up until 810*ms* after the onset of the target word (3-way interaction cluster-based permutation. *p <* 0.05; Maris and Oostenveld 60).

In summary, our results provide the first demonstration of the markedness effect in human neurophysiological data.

## Discussion

Human language processing requires the processing of hierarchical representations in the form of nested tree structures. The existence of such representations in humans is hardly in doubt, since it was demonstrated across a wide range of studies in linguistics, psycholinguistics, and neuroscience. However, online language processing might also involve other, lower-level, processes, unrelated to hierarchical structures and subject to general cognitive constraints.

We focused on long-range subject-verb agreement in language, such as in sentence 1, because this is a domain in which previous behavioral research suggests the involvement of both hierarchical and non-hierarchical processes. The long-range agreement requires hierarchical processing since to determine the correct grammatical number of the verb, one must represent the hierarchical structure of the sentence and use it to neglect other nouns that occur in intermediate prepositional or relative phrases but are not the genuine subject of the verb. For example, in sentences like “The boy near the girl likes climbing,” the brain employs hierarchical processing to determine that it is the ‘boy’, not the ‘girl’, who enjoys climbing. However, a variety of behavioral experiments also indicate that the processing of such sentences can suffer from interference from non-hierarchically linked nouns, such as *N*_2_ in sentence 1 (e.g., 38, 41).

We reasoned, however, that *N*_2_ could interfere linearly in the processing of the verb in various ways, which could lead to similar behavioral patterns. We studied two such, non-mutually exclusive, mechanisms. In the first, a local, sequential mechanism, sensitive to local transition probabilities, *N*_2_ generates a prediction opposite to that of *N*_1_ about the up-coming verb. In the second, the processing of *N*_2_ interferes with the stored representation of *N*_1_, leading to a more fragile or downright erroneous representation of the grammatical number of *N*_1_. Both mechanisms would lead to errors in the processing of the verb, but due to different reasons: one is based on conflicting predictions, the other on memory interference.

Studies on the processing of sequences of non-linguistic stimuli, in both humans and monkeys, have previously supported the first alternative. They showed that there exist two distinct neural mechanisms involved in the processing of non-linguistic sequence patterns, one sensitive to global regularities in the sequence, and the other sensitive only to local transitions between adjacent items (23). Such global and local predictions would be analog to predictions from the main (*N*_1_) and local (*N*_2_) nouns, respectively. Furthermore, analyses of artificial neural networks have concluded that both structure-sensitive and structure-agnostic mechanisms contribute to next-word prediction in AI language models (73). Findings from these models predict that, in humans too, two distinct mechanisms might underlie next-word prediction.

Support for the second alternative comes from numerous studies showing interference effects among words stored in memory (e.g., 74–76). In particular, the cue-based retrieval model (40) suggests that during the processing of the main verb, the local noun *N*_2_ can interfere with the retrieval of the main noun *N*_1_ and cause an erroneous representation of *N*_1_’s features (e.g., its grammatical number). This can in turn lead to subject-verb agreement errors.

In this work, we conducted a neuroimaging experiment using magnetoencephalography (MEG) to study the neural correlates of long-distance agreement during sentence processing and to tease apart these alternatives. We found that contrary to the conflicting prediction signals hypothesis, there was no evidence for a local transition effect. Remarkably, the human brain does not generate any decodable error signal in response to local violations of grammatical number such as “the girls likes”, as long as these violations can be accounted for by long-distance agreement with a distant subject, for instance, “The boy near the girls likes climbing”. The only decodable signals arose from violations of long-distance agreement. This observation is all the more impressive that, in the non-linguistic domain, exactly the converse is found: local violation signals are early and large, while global violation signals are slower and more delayed (31, 33, 77, 78) . Furthermore, our results present positive evidence for the memory-interference hypothesis: the congruity of the two nouns modulated the size of the response to long-distance violations, as predicted by the memory-interference account. We next discuss the implications of these results in view of previous findings from the literature.

## A. Distinct mechanisms for the processing of sentences versus non-linguistic stimuli

The local-global paradigm is a variant of the auditory oddball paradigm (23), in which participants are presented with sounds in short sequences instead of in a continuous manner as in the classic oddball paradigm, thus allowing participants to chunk them and to predict an entire chunk. The local-global paradigm shows that when participants are presented with *aaaab* sequence patterns, in which the first four tones are identical and the fifth differs, the deviancy of the last tone generates a mismatch response (MMR) followed by a late surprise-elicited *P* 3*b* wave (23). A repetition of the same *aaaab* sequence reduces the *P* 3*b* component, however, while the “local” effect of MMR remains, suggesting that the MMR is an automatic response to local transition probabilities. The disappearance of the *P* 3*b* component suggests that a “global” expectation for a deviant fifth tone was generated. Indeed, when a *aaaaa* pattern is subsequently presented, the *P* 3*b* wave reappears, showing that a monotonic sequence can be surprising if it violates prior expectations (79).

The neural signatures of the local and global effects further differ in several ways. First, the local effect is early (100–200ms) and transient, whereas the global effect requires an additional 100-200 ms to rise, and it remains stable (32). Second, the local effect is automatic (does not require attention) and unconscious, whereas the global one disappears when participants are not attending or unconscious (23, 33). Third, while the local effect was traced back to auditory cortices (30, 80), the global one is distributed across the superior temporal sulcus, inferior frontal gyrus, dorsolateral prefrontal, intraparietal, anterior, and posterior cingulate cortices (77). Taken together, these findings show the co-existence of two distinct neural mechanisms involved in the processing of sequence patterns, with very different properties and which are sensitive to regularities at different time scales (34, 35).

In contrast to the processing of auditory stimuli, the results from sentence processing differ substantially. For all three constructions, we found no transition-based effect, and only the structural effect was significant (Figure 4). This suggests that during language processing, structure-based computations dominate over transition-based computations, which is opposite to the case of non-linguistic stimuli. Specifically, in the local-global paradigm with sequences of auditory tones, transition effects are easily detectable, while structural effects can be fragile and get reduced, for instance, when subjects are distracted. For language processing, this is the opposite: while the structural effect is large and easily detectable, the transition effect is barely detectable or nonexistent. Thus, once sequence items enter into the language system, as is the case here for features of number and animacy, sentence-level computations are entirely dominated by structure-sensitive processes, and largely robust to low-level transition effects.

## B. Distinct mechanisms underlying long-range agreement in humans and in language models

Recent studies on neural language models show that structure-sensitive neural mechanisms dedicated to the processing of long-range agreements naturally emerge in the models during training (36). Specifically, it was found that long-range agreements are processed in the models by a small neural circuit, composed of a few specialized units. The core of this neural circuit contains two types of units, termed ‘Syntax units’ and ‘Long-range number units’. The syntax units are sensitive to the latent structure of the sentence, and they convey this information to the number units, which in turn carry the grammatical information of the main subject up to the verb. This mechanism was found to emerge in the models during training without explicit supervision (the models were merely trained on a next-word prediction task), consistently across languages and for various grammatical features (grammatical number and gender) (37).

Furthermore, it was found that grammatical agreement is, in fact, processed by both a structure-sensitive and a structure-agnostic mechanism. The latter is a much simpler mechanism, which processes words independently of the latent hierarchical structure of the sentence (36). This second type of mechanism was shown to be sensitive to mere local word transitions in language and to be carried by a larger set of units, termed ‘Short-range number units’. The short-range units carry predictions about upcoming words based on surface statistical regularities among word sequences, whereas the long-range units generate predictions based on the hierarchical structure of the sentence.

Thus, in language models, predictions about upcoming words arise from two processes occurring at distinct time scales and distinct levels of encoding of incoming sequences. In the present study, we confirmed this dual organization using decoding. For all three constructions tested, we found significant decoding of the structural, transition, and congruity effects (Figure 4). For the structural and transition effects, maximal decoding was reached at the verb onset, whereas for the congruity effect it occurred on the preceding noun, since information about congruity is already available at this point. The presence of both structural and transition effects is in accordance with the predictions from previous analyses, which identified two distinct types of mechanisms in the models.

By contrast, the absence of a transition effect in humans points to a substantial difference between the neural mechanisms underlying long-range agreements in humans and language models. In humans, the structural effect is the dominant one and transition-based predictions were undetectable. In the models, although both effects were equally well decoded, the transition effect corresponds to a larger number of units compared to the structural one. Specifically, the transition effect corresponds to neural activity of the ‘short-range units’, which are abundant in the model, whereas the structural effect corresponds to neural activity of only a small set of units and was shown to be highly ‘sparse’ (37). Behavioral evidence for such sparsity was also recently reported for the newer Transformer architecture (81). In summary, similarly to the case of the processing of non-linguistic stimuli, the structural effect is the dominant one when it comes to human language processing, unlike what occurs in current language models.

## C. Neural correlates of memory interference between the main and embedded nouns during sentence processing

Memory-based models of sentence processing (40, 43, 82–86) suggest that new incoming materials, such as an inflected verb, trigger memory retrieval of previous information, stored in sentence constituents in memory, in order to complete the noun-verb pairing process. The cue-based model is a two-stage mechanism in which the parser predicts the number of the verb and only engages in a retrieval process when this prediction mismatches the bottom-up input. The second stage of this mechanism is sensitive to interference effects and might lead to the retrieval of the wrong feature (e.g., grammatical number).

Note that since predictions about the verb in the cue-based retrieval model depend on a parser, they are only structural, determined by the main noun *N*_1_ only. Low-level sequential predictions, as in the case of sequences of non-linguistic stimuli and neural language models, are not considered in the cue-based retrieval model. Therefore, in memory-based models, a local prediction signal arising from *N*_2_ alone does not occur and is not needed to explain the observed behavioral interference. Our results are fully consistent with this view, without having to assume any additional low-level prediction mechanisms (as present, for instance, in neural language models). First, our behavioral results show a positive interaction between grammaticality and congruity (i.e., a transition effect; Figure 3). That is, an incongruity between the main noun and the local noun elicited more errors in ungrammatical compared to grammatical sentences, more so than in the congruent conditions. This finding is akin to a previous effect reported in studies using self-paced reading, which was termed ‘grammatical asymmetry’ (43, 70). In these studies, it was shown that agreement attraction facilitated the processing of ungrammatical but not grammatical sentences, in the case of incongruent nouns, in accordance with a cue-based retrieval model—a retrieval process was triggered only when the prediction of the verb mismatched the bottom-up input. Our behavioral results, therefore, replicate the ‘grammatical asymmetry’ effect.

Furthermore, in MEG signals, we found that the structural effect was modulated by congruency (Figure 5). This provides positive neural evidence for memory interference between incongruent nouns during agreement processing, in support of memory-interference models. Since a transition effect was not observed in the neural data, memory interference provides a sufficient explanation for agreement errors observed behaviorally and rejects the need to assume that transition-based predictions are active alongside structured predictions. In sum, our results reject the conflicting prediction signal hypothesis and corroborate the memory-interference one.

## D. Discrepancy between the processing of grammatical number and animacy

The behavioral data revealed a large difference between a violation with respect to grammatical number and with respect to animacy. Participants were able to detect both number and animacy violations and, in both cases, were better at affirming that a sentence is grammatical/felicitous than at detecting a violation. However, an intervening incongruent noun, with a mismatching grammatical number, induced a large behavioral interference in the case of number violations, while a similar intervention of a noun with a mismatching animacy feature did not affect participant performance in the case of animacy violations.

Although both number and animacy are syntactic as well as semantic features, number is borne by an overt morpheme in both nouns and verbs, whereas animacy information can only be retrieved from the lexicon. Thus, number may have been processed at a morphosyntactic stage earlier than, and more susceptible to interference than, the lexicose-mantic stage which is needed to detect animacy violations. Our results suggest that the latter stage is totally structure-dependent and immune to intervention by an incongruent noun. At the very least, they indicate that during language processing, grammatical number, and animacy are processed and integrated into an ongoing sentence representation in quite different ways and that the processing of animacy is more robust to intervening material.

This result is, again, consistent with memory-based models of sentence processing (40) for which it was suggested that morphosyntactic processing is relatively ‘fragile’ compared to the processing of animacy (87). Memory-based models of sentence processing suggest that new incoming materials, such as an inflected verb, trigger memory retrieval of previous information, stored in sentence constituents in memory, in order to complete the noun-verb pairing process. This retrieval process is sensitive to similarities among items in memory, and can therefore explain the observed discrepancy. Morphosyntactic features were found to be weaker cues compared to animacy (87), thus resulting in higher similarities among memory items that only differ in morphosyntactic marking. This could make grammatical-number processing more prone to confusion errors, compared to animacy, and therefore to more erroneous grammaticality judgments. Our results therefore corroborate the robustness of animacy processing compared to grammatical number.

## E. General discussion

Our study is subject to several limitations. First, the absence of evidence is not evidence of absence, and the transition effect might have been undetected due to the low signal-to-noise ratio of the MEG data. However, the modulation of the structural effect by congruity (Figure 5A), and further, by grammatical number (Figure 5B&C), shows that our decoding techniques are quite sensitive and highly likely to reflect linguistic processing. This suggests that, even if transition-based computations do occur during sentence processing, their neural traces are small compared to those related to structure-based computations.

Second, our participants were explicitly instructed to detect either grammatical or semantic violations. It is therefore possible that the decoding of neural responses to these violations reflected, at least in part, the imposed task demands rather than pure linguistic processing. Do note that we randomized the assignments of the motor responses within each subject, thus making them orthogonal to violations and therefore unlikely to be at the source of the observed decodings. Still, task demands could have amplified the detection of violations by drawing attention to them. Nevertheless, we see no reason why attention could not have had the additional effect of amplifying a local transition effect if such an effect existed. It is, therefore, reasonable to assume that the core of our conclusions would still hold, albeit with even more reduced and therefore harder-to-detect MEG signals, if we had adopted a passive sentence reading paradigm. In particular, the conclusions regarding the large discrepancy between the structural and transition effects as well as the modulation of the structural effect by congruency, which is orthogonal to such possible amplification, would still hold. A passive task would have also incurred other concerns, for example, the difficulty of participants to engage in the task throughout the experiment, and the relative invisibility of the small morphological markers of agreement in French.

Third, our study is entirely focused on grammatical agreement, and therefore its conclusions are limited to this scope. There might well be transition effects in simpler conditions, and indeed word-transition and syllable-transition probabilities effects have been reported during story listening (88). Our study merely shows that, once the stimuli enter a linguistic processing stage in which the morphological markers for number are involved, then no local transition probability effects are detectable.

Last, we note that our study is not the first to provide neural evidence in favor of the cue-based retrieval model (e.g., 89). However, the unique setup proposed in our study is to directly contrast hierarchical and sequential processing during sentence comprehension, and to further disentangle two possible alternatives for behavioral interference effects.

In conclusion, we summarize the main contributions of our study. First, at the expense of the processing of non-linguistic stimuli and with neural language models, sentence processing in the human brain seems largely robust to low-level transition-based, predictions. The present study provides neural evidence in favor of the dominance of structure-based over transition-based processing during sentence processing. Second, we showed positive neural evidence for memory interference during sentence processing, which provides support for cue-based retrieval models in explaining long-range subject-verb agreement errors. Last, we provide, to our knowledge, the first neural evidence for the markedness effect in sentence processing.

## ACKNOWLEDGEMENTS

The authors would like to thank Christophe Pallier, Théo Desbordes, JeanRémi King, and Luigi Rizzi for their interesting discussions and feedback on this work. We are also grateful to Fosca Al Roumi, Leila Azizi, as well as the MEG team of ICM for helping with the data acquisition. This work was supported by INSERM, CEA, Collège de France, the Bettencourt-Schueller Foundation, and an ERC grant, the “NeuroSyntax” project, to S.D.

## Data and Code Availability

The data and code supporting this study are available upon request.

## Author Contributions

- **C.Z:** Methodology, Writing, Conceptualization, Data Collection, Data Analysis.
- **S.D:** Conceptualization, Methodology, Writing, Funding, Project Administration, Validation.
- **Y.L:** Conceptualization, -0Methodology, Writing, Project Administration, Validation.

## Supplementary Note A: Lexicon

### Nouns

#### Singular

boy, father, husband, brother, man, nephew, girl, mother, wife, sister, woman, niece, athlete, baker, doctor, farmer, teacher, lawyer, actor, author, banker, barber, chef, dentist, judge, nurse, painter, pilot, plumber, tailor, waiter, vet, builder, car, bus, taxi, truck, tractor, scooter, van

#### Plural

boys, fathers, husbands, brothers, men, nephews, girls, mothers, wives, sisters, women, nieces, athletes, bakers, doctors, farmers, teachers, lawyers, actors, authors, bankers, barbers, chefs, dentists, judges, nurses, painters, pilots, plumbers, tailors, waiters, vets, builders, cars, buses, taxis, trucks, tractors, scooters, vans

### Verbs

#### Singular

likes, loves, hates, avoids, dislikes, fears, prefers, detests, departs, moves, turns, stops, prays, sits, dies, arrives, leaks, rusts, malfunctions

#### Plural

like, love, hate, avoid, dislike, fear, prefer, detest, depart, move, turn, stop, pray, sit, die, arrive, leak, rust, malfunction

### Adverbs

abruptly, suddenly

### Activities

climbing, skiing, cooking, shopping, painting, studying, walking, cycling, farming, fencing, gambling, knitting, acting, boxing, bowling, camping, fishing, skating, dancing, sailing, yachting, hunting, spinning, driving

### Prepositions

near, by, beside

### Pronouns

that

## Supplementary Note B: Instructions

## Supplementary Note C: Stimuli from the first run of the experiment

## Supplementary Note D: Effect of response type on the neural data

## Footnotes

1 https://dialangweb.lancaster.ac.uk/

2 The lexicon used in the study is presented in appendix A.

3 The exact instructions are presented in Appendix B

4 https://github.com/facebookresearch/colorlessgreenRNNs

## Bibliography

1. Noam Chomsky. Syntactic structures. Mouton publishers, 1957. ISBN 978-90-279-3385-0. Accepted: 2019-04-10T06:25:21Z Journal Abbreviation: Janua linguarum.

2. Marc Hauser, Noam Chomsky, and Tecumseh Fitch. The faculty of language: What is it, who has it, and how did it evolve? Science, 298(5598):1569–1579, 11 2002.

3. Martin BH Everaert, Marinus AC Huybregts, Noam Chomsky, Robert C Berwick, and Johan J Bolhuis. Structures, not strings: linguistics as part of the cognitive sciences. Trends in cognitive sciences, 19(12):729–743, 2015.

4. Michiru Makuuchi, Jörg Bahlmann, Alfred Anwander, and Angela D Friederici. Segregating the core computational faculty of human language from working memory. Proceedings of the National Academy of Sciences, 106(20):8362–8367, 2009.

5. Stefan Frank, Rens Bod, and Morten Christiansen. How hierarchical is language use? Proceedings of the Royal Society B: Biological Sciences, 279(1747):4522–4531, 2012.

6. Ewa Da?browska and Elena Lieven. Towards a lexically specific grammar of children’s question constructions. 2005.

7. Ewa Da?browska. What exactly is universal grammar, and has anyone seen it? Frontiers in psychology, 6:852, 2015.

8. Noam Chomsky. Lectures on government and binding: The Pisa lectures. Number 9. Walter de Gruyter, 1993.

9. Noam Chomsky, Peter W Culicover, Thomas Wasow, Adrian Akmajian, et al. On wh-movement. 1977, 65, 1977.

10. Noam Chomsky. Problems of knowledge and freedom: The russell lectures. 1971.

11. Massimo Piattelli-Palmarini. Language and learning: the debate between jean piaget and noam chomsky. 1980.

12. Mark Liberman and Alan Prince. On stress and linguistic rhythm. Linguistic inquiry, 8(2): 249–336, 1977.

13. Noam Chomsky. Essays on Form and Interpretation. North Holland, New York, 1977.

14. Joseph Aoun and David W Lightfoot. Government and contraction. Linguistic Inquiry, 15 (3):465–473, 1984.

15. Mariacristina Musso, Andrea Moro, Volkmar Glauche, Michel Rijntjes, Jürgen Reichenbach, Christian Büchel, and Cornelius Weiller. Broca’s area and the language instinct. Nature neuroscience, 6(7):774–781, 2003.

16. Michal Ben-Shachar, Dafna Palti, and Yosef Grodzinsky. Neural correlates of syntactic movement: converging evidence from two fmri experiments. Neuroimage, 21(4):1320–1336, 2004.

17. Angela D Friederici, Jörg Bahlmann, Stefan Heim, Ricarda I Schubotz, and Alfred Anwander. The brain differentiates human and non-human grammars: functional localization and structural connectivity. Proceedings of the National Academy of Sciences, 103(7):2458–2463, 2006.

18. Angela D Friederici. The brain basis of language processing: from structure to function. Physiological reviews, 91(4):1357–1392, 2011.

19. Christophe Pallier, Anne-Dominique Devauchelle, and Stanislas Dehaene. Cortical representation of the constituent structure of sentences. Proceedings of the National Academy of Sciences, 108(6):2522–2527, 2011.

20. Einat Shetreet and Naama Friedmann. The processing of different syntactic structures: fmri investigation of the linguistic distinction between wh-movement and verb movement. Journal of Neurolinguistics, 27(1):1–17, 2014.

21. Emiliano Zaccarella and Angela D Friederici. Merge in the human brain: A sub-region based functional investigation in the left pars opercularis. Frontiers in psychology, 6:1818, 2015.

22. Matthew Nelson, Imen El Karoui, Kristof Giber, Xiaofang Yang, Laurent Cohen, Hilda Koopman, Sydney Cash, Lionel Naccache, John Hale, Christophe Pallier, and Stanislas Dehaene. Neurophysiological dynamics of phrase-structure building during sentence processing. Proceedings of the National Academy of Sciences, 114(18):E3669–E3678, 2017.

23. Tristan A Bekinschtein, Stanislas Dehaene, Benjamin Rohaut, François Tadel, Laurent Cohen, and Lionel Naccache. Neural signature of the conscious processing of auditory regularities. Proceedings of the National Academy of Sciences, 106(5):1672–1677, 2009.

24. Stefan Koelsch, Martin Rohrmeier, Renzo Torrecuso, and Sebastian Jentschke. Processing of hierarchical syntactic structure in music. Proceedings of the National Academy of Sciences, 110(38):15443–15448, 2013.

25. Stefan Koelsch, Tobias Busch, Sebastian Jentschke, and Martin Rohrmeier. Under the hood of statistical learning: A statistical mmn reflects the magnitude of transitional probabilities in auditory sequences. Scientific reports, 6(1):19741, 2016.

26. Vincent KM Cheung, Lars Meyer, Angela D Friederici, and Stefan Koelsch. The right inferior frontal gyrus processes nested non-local dependencies in music. Scientific reports, 8(1): 3822, 2018.

27. Vera Tsogli, Sebastian Jentschke, Tatsuya Daikoku, and Stefan Koelsch. When the statistical mmn meets the physical mmn. Scientific reports, 9(1):5563, 2019.

28. Vera Tsogli, Sebastian Jentschke, and Stefan Koelsch. Unpredictability of the “when” influences prediction error processing of the “what” and “where”. PloS one, 17(2):e0263373, 2022.

29. Vera Tsogli, Stavros Skouras, and Stefan Koelsch. Brain-correlates of processing local dependencies within a statistical learning paradigm. Scientific reports, 12(1):15296, 2022.

30. Imen El Karoui, Jean-Remi King, Jacobo Sitt, Florent Meyniel, Simon Van Gaal, Dominique Hasboun, Claude Adam, Vincent Navarro, Michel Baulac, Stanislas Dehaene, et al. Event-related potential, time-frequency, and functional connectivity facets of local and global auditory novelty processing: an intracranial study in humans. Cerebral cortex, 25(11):4203–4212, 2015.

31. Zenas C Chao, Kana Takaura, Liping Wang, Naotaka Fujii, and Stanislas Dehaene. Large-scale cortical networks for hierarchical prediction and prediction error in the primate brain. Neuron, 100(5):1252–1266, 2018.

32. Jean-Rémi King and Stanislas Dehaene. Characterizing the dynamics of mental representations: the temporal generalization method. Trends in cognitive sciences, 18(4):203–210, 2014.

33. Melanie Strauss, Jacobo D Sitt, Jean-Remi King, Maxime Elbaz, Leila Azizi, Marco Buiatti, Lionel Naccache, Virginie Van Wassenhove, and Stanislas Dehaene. Disruption of hierarchical predictive coding during sleep. Proceedings of the National Academy of Sciences, 112(11):E1353–E1362, 2015.

34. Florent Meyniel, Maxime Maheu, and Stanislas Dehaene. Human inferences about sequences: A minimal transition probability model. PLoS computational biology, 12(12): e1005260, 2016.

35. Maxime Maheu, Stanislas Dehaene, and Florent Meyniel. Brain signatures of a multiscale process of sequence learning in humans. elife, 8:e41541, 2019.

36. Yair Lakretz, German Kruszewski, Theo Desbordes, Dieuwke Hupkes, Stanislas Dehaene, and Marco Baroni. The emergence of number and syntax units in lstm language models, 2019.

37. Yair Lakretz, Dieuwke Hupkes, Alessandra Vergallito, Marco Marelli, Marco Baroni, and Stanislas Dehaene. Mechanisms for handling nested dependencies in neural-network language models and humans. Cognition, 213:104699, 2021. ISSN 0010-0277. doi: 10.1016/j.cognition.2021.104699.

38. Kathryn Bock and Carol A Miller. Broken agreement. Cognitive psychology, 23(1):45–93, 1991.

39. Robert J Hartsuiker, Inés Antón-Méndez, and Marije Van Zee. Object attraction in subject-verb agreement construction. Journal of Memory and Language, 45(4):546–572, 2001.

40. Richard L Lewis and Shravan Vasishth. An activation-based model of sentence processing as skilled memory retrieval. Cognitive science, 29(3):375–419, 2005.

41. Julie Franck, Glenda Lassi, Ulrich H Frauenfelder, and Luigi Rizzi. Agreement and movement: A syntactic analysis of attraction. Cognition, 101(1):173–216, 2006.

42. Julie Franck, Ulrich Hans Frauenfelder, and Luigi Rizzi. A syntactic analysis of interference in subject–verb agreement. MIT working papers in linguistics, (53):173–190, 2007.

43. Matthew W Wagers, Ellen F Lau, and Colin Phillips. Agreement attraction in comprehension: Representations and processes. Journal of Memory and Language, 61(2):206–237, 2009.

44. Nicola Molinaro, Horacio A Barber, and Manuel Carreiras. Grammatical agreement processing in reading: Erp findings and future directions. cortex, 47(8):908–930, 2011.

45. Nicola Molinaro, Francesco Vespignani, Roberto Zamparelli, and Remo Job. Why brother and sister are not just siblings: Repair processes in agreement computation. Journal of Memory and Language, 64(3):211–232, 2011.

46. Lee Osterhout and Phillip J Holcomb. Event-related brain potentials elicited by syntactic anomaly. Journal of memory and language, 31(6):785–806, 1992.

47. Lee Osterhout and Linda A Mobley. Event-related brain potentials elicited by failure to agree. Journal of Memory and language, 34(6):739–773, 1995.

48. Karl Friston. A theory of cortical responses. 360(1456):815–836, April 2005. doi: 10.1098/rstb.2005.1622.

49. Gina R Kuperberg and T Florian Jaeger. What do we mean by prediction in language comprehension? Language, cognition and neuroscience, 31(1):32–59, 2016.

50. Karl Friston, Rosalyn J. Moran, Yukie Nagai, Tadahiro Taniguchi, Hiroaki Gomi, and Josh Tenenbaum. World model learning and inference. 144:573–590, December 2021. doi: 10.1016/j.neunet.2021.09.011.

51. Angela D Friederici, Karsten Steinhauer, and Stefan Frisch. Lexical integration: Sequential effects of syntactic and semantic information. Memory & cognition, 27(3):438–453, 1999.

52. Peter Hagoort. Interplay between syntax and semantics during sentence comprehension: Erp effects of combining syntactic and semantic violations. Journal of cognitive neuroscience, 15(6):883–899, 2003.

53. Alexandre Gramfort, Martin Luessi, Eric Larson, Denis A Engemann, Daniel Strohmeier, Christian Brodbeck, Roman Goj, Mainak Jas, Teon Brooks, Lauri Parkkonen, et al. Meg and eeg data analysis with mne-python. Frontiers in neuroscience, 7:267, 2013.

54. Mainak Jas, Eric Larson, Denis A Engemann, Jaakko Leppäkangas, Samu Taulu, Matti Hämäläinen, and Alexandre Gramfort. A reproducible meg/eeg group study with the mne software: recommendations, quality assessments, and good practices. Frontiers in neuroscience, 12:530, 2018.

55. Samu Taulu, Matti Kajola, and Juha Simola. Suppression of interference and artifacts by the signal space separation method. Brain topography, 16(4):269–275, 2004.

56. François Perrin, Jacques Pernier, O Bertrand, and Jean Francois Echallier. Spherical splines for scalp potential and current density mapping. Electroencephalography and clinical neurophysiology, 72(2):184–187, 1989.

57. Stanislas Dehaene and Jean-Rémi King. Decoding the dynamics of conscious perception: The temporal generalization method. Micro-, meso-and macro-dynamics of the brain, pages 85–97, 2016.

58. Fabian Pedregosa, Gaël Varoquaux, Alexandre Gramfort, Vincent Michel, Bertrand Thirion, Olivier Grisel, Mathieu Blondel, Peter Prettenhofer, Ron Weiss, Vincent Dubourg, et al. Scikit-learn: Machine learning in python. the Journal of machine Learning research, 12: 2825–2830, 2011.

59. D Bates, M Maechler, B Bolker, and S Walker. lme4: Linear mixed-effects models using eigen and s4. r package version 1.1–7. 2014, 2015.

60. Eric Maris and Robert Oostenveld. Nonparametric statistical testing of eeg-and meg-data. Journal of neuroscience methods, 164(1):177–190, 2007.

61. Yoav Goldberg. Neural Network Methods for Natural Language Processing. Morgan & Claypool, San Francisco, CA, 2017.

62. Kristina Gulordava, Piotr Bojanowski, Edouard Grave, Tal Linzen, and Marco Baroni. Colorless green recurrent networks dream hierarchically. In Proceedings of NAACL, pages 1195–1205, New Orleans, LA, 2018.

63. Alex Graves. Supervised Sequence Labelling with Recurrent Neural Networks. Springer, Berlin, 2012.

64. Sepp Hochreiter and Jürgen Schmidhuber. Long short-term memory. Neural Computation, 9(8):1735–1780, 1997.

65. Ofir Press and Lior Wolf. Using the output embedding to improve language models. arXiv preprint arXiv:1608.05859, 2016.

66. Colin Phillips, Matthew W Wagers, and Ellen F Lau. Grammatical illusions and selective fallibility in real-time language comprehension. Experiments at the Interfaces, 37:147–180, 2011.

67. Kathleen M Eberhard. The marked effect of number on subject–verb agreement. Journal of Memory and language, 36(2):147–164, 1997.

68. Gabriella Vigliocco, Brian Butterworth, and Carlo Semenza. Constructing subject-verb agreement in speech: The role of semantic and morphological factors. Journal of Memory and Language, 34(2):186–215, 1995.

69. Kathryn Bock, Manuel Carreiras, and Enrique Meseguer. Number meaning and number grammar in english and spanish. Journal of Memory and Language, 66(1):17–37, 2012.

70. Sol Lago, Diego E Shalom, Mariano Sigman, Ellen F Lau, and Colin Phillips. Agreement attraction in spanish comprehension. Journal of Memory and Language, 82:133–149, 2015.

71. Julie Franck, Gabriella Vigliocco, and Janet Nicol. Subject-verb agreement errors in french and english: The role of syntactic hierarchy. Language and cognitive processes, 17(4): 371–404, 2002.

72. Heidi Lorimor, Kathryn Bock, Ekaterina Zalkind, Alina Sheyman, and Robert Beard. Agreement and attraction in russian. Language and cognitive processes, 23(6):769–799, 2008.

73. Yair Lakretz, Stanislas Dehaene, and Jean-Rémi King. What limits our capacity to process nested long-range dependencies in sentence comprehension? Entropy, 22(4):446, 2020.

74. Lena A Jäger, Felix Engelmann, and Shravan Vasishth. Similarity-based interference in sentence comprehension: Literature review and bayesian meta-analysis. Journal of Memory and Language, 94:316–339, 2017.

75. Richard L Lewis, Shravan Vasishth, and Julie A Van Dyke. Computational principles of working memory in sentence comprehension. Trends in cognitive sciences, 10(10):447–454, 2006.

76. Julie A Van Dyke and Brian McElree. Retrieval interference in sentence comprehension. Journal of memory and language, 55(2):157–166, 2006.

77. Lynn Uhrig, Stanislas Dehaene, and Béchir Jarraya. A hierarchy of responses to auditory regularities in the macaque brain. Journal of Neuroscience, 34(4):1127–1132, 2014.

78. Liping Wang, Lynn Uhrig, Bechir Jarraya, and Stanislas Dehaene. Representation of numerical and sequential patterns in macaque and human brains. Current Biology, 25(15): 1966–1974, 2015.

79. Stanislas Dehaene, Florent Meyniel, Catherine Wacongne, Liping Wang, and Christophe Pallier. The neural representation of sequences: from transition probabilities to algebraic patterns and linguistic trees. Neuron, 88(1):2–19, 2015.

80. Felipe Pegado, Tristan Bekinschtein, Nicolas Chausson, Stanislas Dehaene, Laurent Cohen, and Lionel Naccache. Probing the lifetimes of auditory novelty detection processes. Neuropsychologia, 48(10):3145–3154, 2010.

81. Yair Lakretz, Théo Desbordes, Dieuwke Hupkes, and Stanislas Dehaene. Can transformers process recursive nested constructions, like humans? In Proceedings of the 29th International Conference on Computational Linguistics, pages 3226–3232, Gyeongju, Republic of Korea, October 2022. International Committee on Computational Linguistics.

82. Brian McElree, Stephani Foraker, and Lisbeth Dyer. Memory structures that subserve sentence comprehension. Journal of memory and language, 48(1):67–91, 2003.

83. Julie A Van Dyke and Clinton L Johns. Memory interference as a determinant of language comprehension. Language and linguistics compass, 6(4):193–211, 2012.

84. William Badecker and Frantisek Kuminiak. Morphology, agreement and working memory retrieval in sentence production: Evidence from gender and case in slovak. Journal of memory and language, 56(1):65–85, 2007.

85. Brian Dillon, Alan Mishler, Shayne Sloggett, and Colin Phillips. Contrasting intrusion profiles for agreement and anaphora: Experimental and modeling evidence. Journal of Memory and Language, 69(2):85–103, 2013.

86. Andrea E Martin and Brian McElree. A content-addressable pointer mechanism underlies comprehension of verb-phrase ellipsis. Journal of Memory and Language, 58(3):879–906, 2008.

87. Anastasia Stoops and Kiel Christianson. Parafoveal processing of inflectional morphology on russian nouns. Journal of Cognitive Psychology, 29(6):653–669, 2017.

88. Micha Heilbron, Kristijan Armeni, Jan-Mathijs Schoffelen, Peter Hagoort, and Floris P de Lange. A hierarchy of linguistic predictions during natural language comprehension. bioRxiv, pages 2020–12, 2021.

89. Andrea E Martin, Mante S Nieuwland, and Manuel Carreiras. Event-related brain potentials index cue-based retrieval interference during sentence comprehension. NeuroImage, 59 (2):1859–1869, 2012.

